# A dynamic sequence of visual processing initiated by gaze shifts

**DOI:** 10.1101/2022.08.23.504847

**Authors:** Philip R. L. Parker, Dylan M. Martins, Emmalyn S. P. Leonard, Nathan M. Casey, Shelby L. Sharp, Elliott T. T. Abe, Matthew C. Smear, Jacob L. Yates, Jude F. Mitchell, Cristopher M. Niell

## Abstract

Animals move their head and eyes as they explore and sample the visual scene. Previous studies have demonstrated neural correlates of head and eye movements in rodent primary visual cortex (V1), but the sources and computational roles of these signals are unclear. We addressed this by combining measurement of head and eye movements with high density neural recordings in freely moving mice. V1 neurons responded primarily to gaze shifts, where head movements are accompanied by saccadic eye movements, but not to head movements where compensatory eye movements stabilize gaze. A variety of activity patterns immediately followed gaze shifts, including units with positive, biphasic, or negative responses, and together these responses formed a temporal sequence following the gaze shift. These responses were greatly diminished in the dark for the vast majority of units, replaced by a uniform suppression of activity, and were similar to those evoked by sequentially flashed stimuli in head-fixed conditions, suggesting that gaze shift transients represent the temporal response to the rapid onset of new visual input. Notably, neurons responded in a sequence that matches their spatial frequency preference, from low to high spatial frequency tuning, consistent with coarse-to-fine processing of the visual scene following each gaze shift. Recordings in foveal V1 of freely gazing head-fixed marmosets revealed a similar sequence of temporal response following a saccade, as well as the progression of spatial frequency tuning. Together, our results demonstrate that active vision in both mice and marmosets consists of a dynamic temporal sequence of neural activity associated with visual sampling.

**Highlights:**

- During free movement, neurons in mouse V1 respond to head movements that are accompanied by a gaze-shifting saccadic eye movement, but not a compensatory eye movement.
- Neurons respond to gaze shifts with diverse temporal dynamics that form a sequence across the population, from early positive responses to biphasic and negative responses.
- In darkness, most neurons show a uniform suppression following a gaze shift.
- Temporal dynamics of responses correspond to a neuron’s temporal and spatial frequency preferences, consistent with a coarse-to-fine processing sequence.
- A similar temporal sequence following saccades is observed in foveal V1 of freely gazing head-fixed marmosets, demonstrating shared aspects of active visual processing across species.

## Introduction

Visual perception is an active process in which animals move through their environment and sample the visual scene through movements of the body, head, and eyes. Self-motion and sampling behavior have a profound impact on the dynamic pattern of retinal visual input, which can play an integral role in processing and interpreting the visual scene (Ahissar and Arieli, 2001; Boi et al., 2017; Gibson, 1979; Schroeder et al., 2010). Correspondingly, a variety of movement-related signals have been observed throughout the brain (Parker et al., 2020). In the visual cortex, eye movement signals have been studied in nonhuman primates and cats (Burchfiel and Duffy, 1974; Leopold and Logothetis, 1998; Nishimoto et al., 2017; Noda and Adey, 1974; Sommer and Wurtz, 2008), and recently in mice (Miura and Scanziani, 2021). Cortex-wide signals of locomotion and body/facial movements are present in head-fixed mice (Musall et al., 2019; Niell and Stryker, 2010; Stringer et al., 2019), and head rotation signals are seen in the rodent visual cortex (Bouvier et al., 2020; Guitchounts et al., 2020; Meyer et al., 2018; Vélez-Fort et al., 2018). Importantly, these signals can be variously attributed to either the movement itself, the consequence of movement on the visual input, or a combination thereof.

However, the computational role of head/eye movement signals in natural vision remains enigmatic, particularly since they are often studied under constrained conditions of head or gaze fixation, with sparse, artificial visual stimuli. Likewise, it is not clear whether there are common principles of active visual processing across rodents and primates, given the differences in their eye movement repertoires and other visual specializations. Indeed, it has recently been shown that the gain modulation of mouse V1 neurons by locomotion (Niell and Stryker, 2010) is present in primates, but differs significantly in magnitude as well as sign (Liska et al., 2022). We therefore sought to characterize the neural representation of visual sampling movements in mice and marmosets, under natural conditions of free movement and viewing.

By measuring both head and eye movements together with neural activity in freely moving mice, we found that most V1 neurons do not respond to compensatory head/eye movements, but respond to gaze-shifting movements with a variety of response dynamics that results in a temporal sequence of population activity following a gaze shift. Under head fixation, when presented with sequentially flashed full-field stimuli, single neuron responses closely matched the neuron’s response to gaze shifts, suggesting that the pattern of gaze shift transients is largely explained by the response to rapid onset of new visual input. The time course of responses corresponded predictably with tuning to increasing spatial frequency, consistent with the coarse-to-fine theory of visual processing (Hegdé, 2008). Notably, we observed a similar sequence of processing following saccades in head-fixed marmosets that were allowed to freely gaze while viewing natural scenes. These results provide new insight into neural signals previously observed around head and eye movements, and reveal a computational principle for coarse-to-fine visual processing during active vision in complex natural scenes that is shared across rodents and primates.

## Results

### Neurons in mouse V1 preferentially respond to gaze-shifting eye/head movements

First, we sought to characterize the responses of individual neurons in mouse V1 to all large-amplitude head and eye movements. To monitor neural responses to head and eye movements during free movement, we used a recording system (Figure 1A) consisting of a head-mounted camera to record the position of the right eye, a forward-facing wide-angle (120 deg) camera to capture the visual scene, an IMU to record head orientation and acceleration, and a chronically-implanted 64- or 128-channel linear silicon probe in primary visual cortex (V1) for single-unit electrophysiology in freely moving animals (Figure 1B; (Michaiel et al., 2020; Parker et al., 2022).

**Figure 1.**
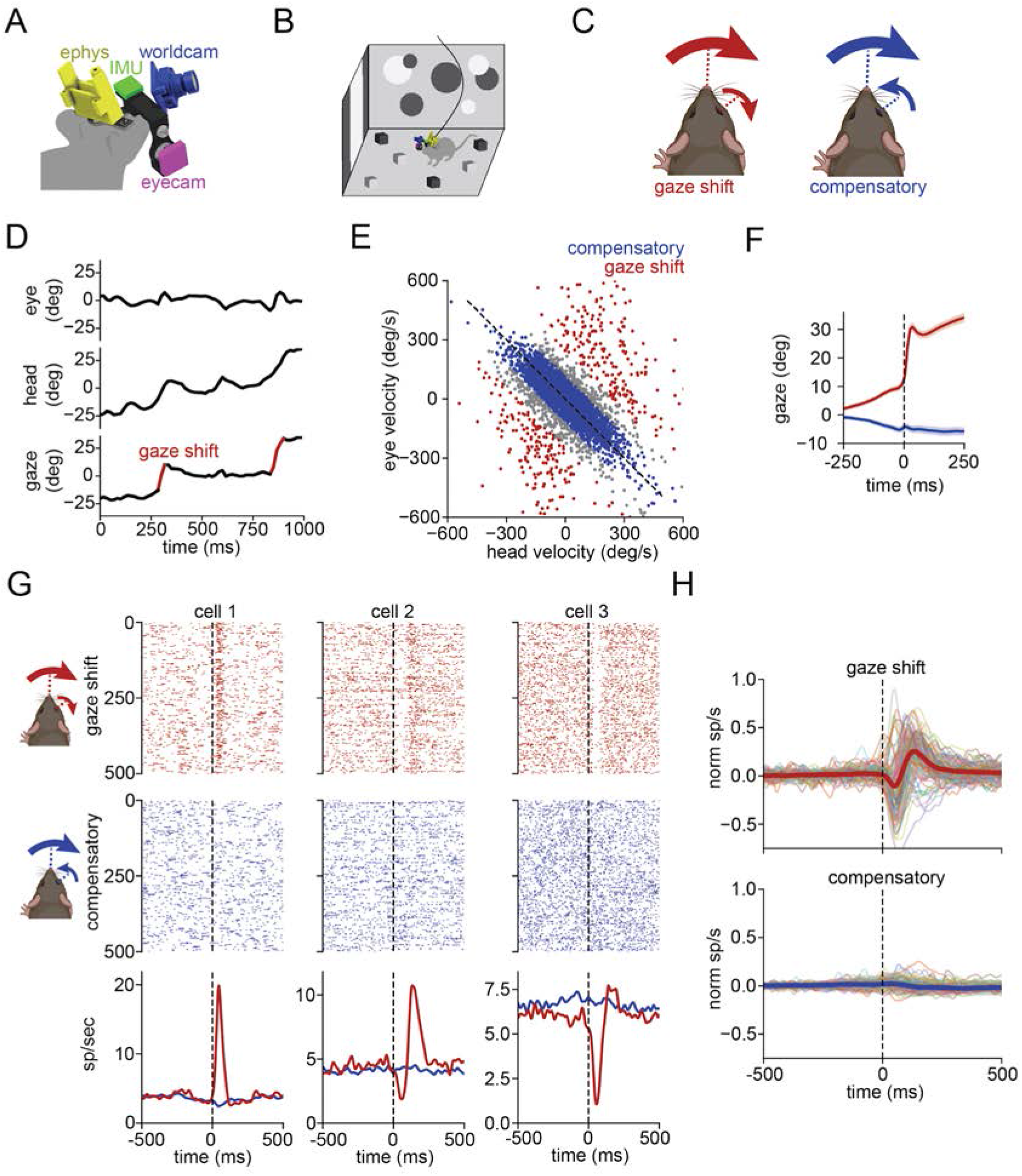
V1 neurons preferentially respond to gaze-shifting eye/head movements. (A) Schematic of the head-mounted recording system. (B) Experimental paradigm, in which a mouse is allowed to freely move through aisually complex environment. (C) Schematic of gaze-shifting and compensatory eye/head movements, in which the eyes either move with or against the movement of the head. (D) Mice change their horizontal eye position (top) to compensate for changing horizontal head position (middle), punctuated by rapid gaze-shifting eye movements. The resulting gaze position (bottom), the sum of horizontal eye and head positions, shows a “saccade-and-fixate” pattern. (E) Scatter plot of eye and head velocities subsampled (25x) from an example 64 min recording, showing segregation of compensatory versus gaze-shifting eye movements. (F) Mean position of the animal’s gaze around gaze-shifting (red) and compensatory (blue) movements in the leftward head direction across all recordings. (G) Response of three example cells to 500 randomly sampled gaze-shifting (top) or compensatory (middle) eye/head movements as a spike raster. Bottom: gaze-shifting and compensatory PSTH. (H) Normalized PSTH of gaze-shifting (top) and compensatory (bottom) eye/head movements for 100 example units with a baseline firing rate >2Hz, with median of all cells (n=716) overlaid.

Mice coordinate their head and eye movements in a “saccade-and-fixate” pattern of selecting and stabilizing the input to the retina that is observed across the animal kingdom (Land, 2019). When the mouse moves its head, the accompanying eye movements can be generally categorized into one of two functional types: compensatory and gaze-shifting (Meyer et al., 2020; Michaiel et al., 2020). Gaze refers to the direction the eyes are pointing relative to the visual scene, i.e. the sum of head direction and eye position within the head, which determines the retinal input. Compensatory eye movements counteract head movements of the opposite direction in order to keep gaze stabilized during self-motion (through vestibulo-ocular and other reflexes), and are typically smooth in trajectory. Conversely, gaze-shifting eye movements accompany head movements in the same direction to change the direction of the animal’s gaze, and are typically abrupt, saccadic movements. Together these lead to a pattern of abrupt shifts of gaze (saccade) interspersed by periods of stabilized gaze (fixate) during continuous head movement.

As observed previously, large amplitude horizontal head movements in our recordings were almost always accompanied by a horizontal eye movement (Meyer et al., 2020; Michaiel et al., 2020). A concurrent eye/head movement when the head and eye move in the same direction, which would shift the direction of gaze, was termed gaze-shifting (Figure 1C, left; 63 ± 11 per min; 3285 ± 893 per experiment over 50 ± 8 min; n=9 mice). Conversely, an eye/head movement when a large eye movement occurs in the opposite direction as the head movement, which would generate no net change in the gaze direction, was termed compensatory (Figure 1C, right; 242 ± 36 per min; 12657 ± 3436 per experiment). The coordinated impact of these two types of movements can be observed by summing together the eye and head position to give the direction of gaze, and reveals the typical saccade-and-fixate pattern with periods of gaze stabilization (fixations) interspersed by abrupt shifts (saccades; Figure 1D). This head movement categorization can be visualized in a plot of eye versus head velocity (Figure 1E), where compensatory movements fall along the negative diagonal, since head and eyes move equal but oppositely, while gaze shifts form a distinct off-diagonal pattern. Based on this, we defined gaze-shifting movements as timepoints with gaze velocity >240 deg/sec (red points in Fig 1E), and compensatory as timepoints <120 deg/sec (blue points in Fig 1E). Aligning gaze position to the time of gaze-shifting or compensatory movements reveals the characteristic step with gaze shifts and little change in gaze position for compensatory movements (Figure 1F).

We extracted the single-unit neural activity in freely moving mice around the time of each gaze-shifting or compensatory eye/head movement (Figure 1G). This revealed striking responses following gaze-shifting movements, including positive, biphasic, and negative responses. Notably, the same modulation is not found around the time of compensatory movements. Across a population of 716 neurons in 9 mice, responses to gaze shifts were abundant (74.2%; 531/716) but compensatory responses were not (6.7%; 48/716; Figure 1H), defined as change in firing of 1 sp/sec and 10% of baseline. Thus, V1 neurons predominantly respond to gaze-shifting eye/head movements, which produce abrupt changes in the retinal input, and do not generally respond to compensatory movements, which stabilize the retinal input. These findings lend support to the notion that some previously observed responses in V1 to head movements may result from the accompanying shift in gaze, rather than the head movement itself.

### Diversity and temporal sequence of gaze-shift responses

Neural responses to gaze shifts were diverse, both in terms of the sign of modulation and the temporal dynamics. In order to characterize this diversity, we first performed k-means clustering of units based on principal component analysis (PCA) of the normalized PSTHs for gaze shifts (Figure S1A). For some units, the response for one direction of horizontal movement was stronger than the opposite direction (Figure S1B). We therefore determined their preferred direction as the direction of movement yielding the largest peak response, and used this PSTH for clustering. This clustering provides a means to describe the diversity within the population, but it is important to note that this is subdividing a continuous population. We selected a value of k = 5 clusters as it captured the diversity of responses while still providing distinct gaze shift motifs in each cluster.

This resulted in four responsive clusters (Figure 2A, 2B top), which showed early positive (11.5% of population; Figure S1C), late positive (18.9%), biphasic (23.7%), and negative (9.2%) responses to gaze shifts, as well as a largely unresponsive fifth cluster (36.7%). None of the clusters had a consistent response to compensatory movements (Figure 2B, bottom). Mean firing rates and the relative fraction of putative excitatory/inhibitory cells based on spike waveform were similar for each cluster (Figure S1D-E). Intriguingly, while most clusters were fairly uniformly distributed in cortical depth, neurons in the early positive population were more likely to be located in the deep layers (Figure S1F). These clusters also corresponded in a predictable manner to the tuning for angular velocity of horizontal head rotation (Figure S1G). Early positive neurons were generally positively tuned for angular velocity, and often selective for one direction of rotation, while biphasic and negative clusters showed negative tuning, consistent with their response immediately after a gaze shift.

**Figure 2.**
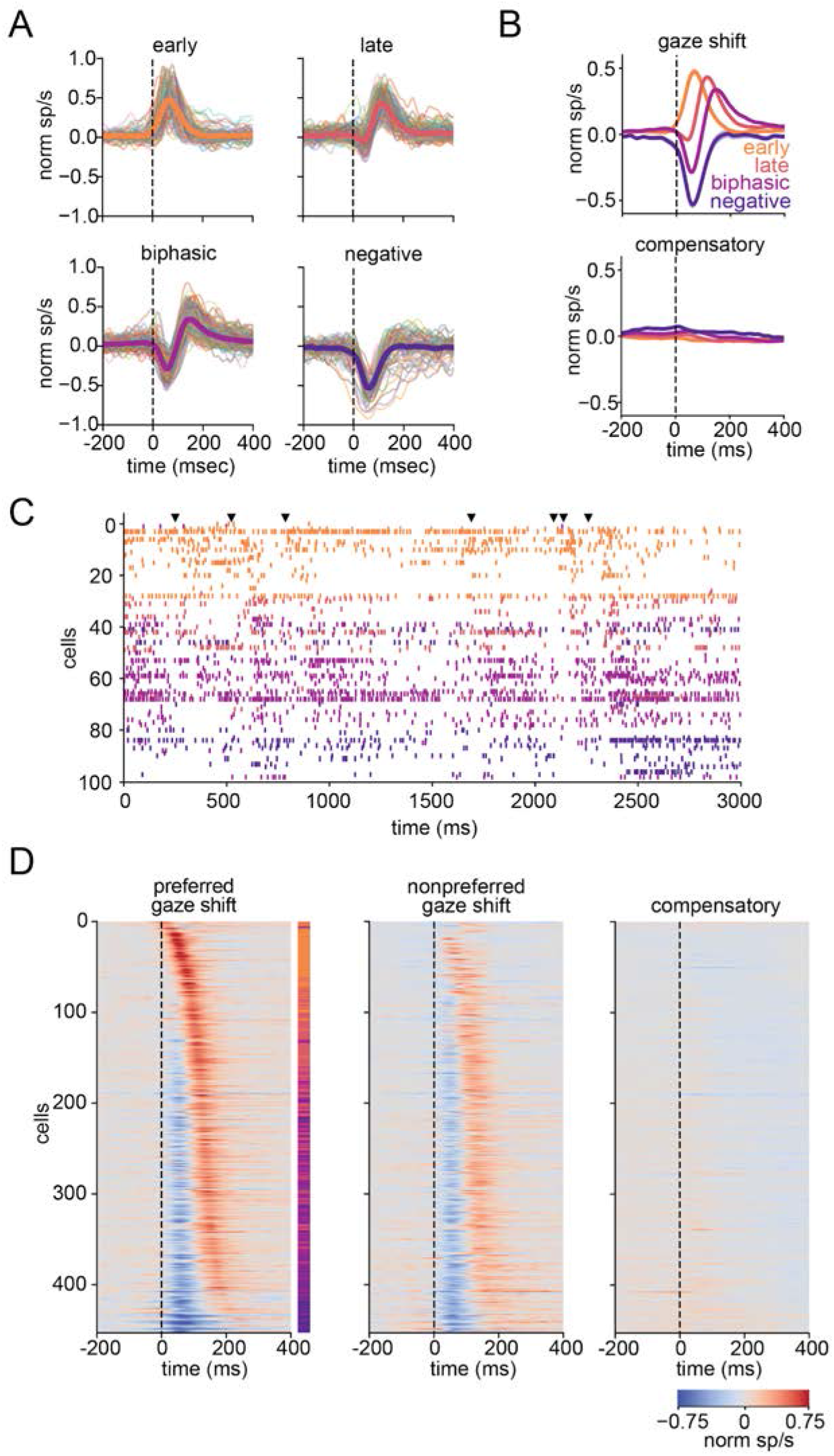
Diversity and temporal sequence of gaze shift responses. (A) Normalized PSTHs for preferred direction of gaze shifts in responsive units, grouped by k-means clustering, with each cluster’s mean overlaid. n=9 mice, 716 units (early=82, late=135, biphasic=170, negative=66, unresponsive=263). (B) Mean normalized PSTH in response to gaze-shifting (top) and compensatory (bottom) eye/head movements, with cells clustered on their gaze shift response. (C) Example spike rasters from a 3 s period of freely moving activity, showing all units responsive to gaze shifts (n=98/128), color coded by response type cluster. Gaze shift onsets are indicated by black arrows (top). (D) Normalized PSTH for gaze-shifting movements in the preferred (left), nonpreferred (middle) and compensatory movements (right), for all responsive cells, sorted along the y-axis by response latency. Vertical color code to the side of the left panel shows each cell’s assigned cluster using the colors in (A).

Notably, the mean traces for each cluster (Figure 2B, top) and the projection in principal component space (Figure S1A) suggested a temporal sequence, from the early positive through the negative. Sorting units based on the time of their peak positive response to gaze shifts revealed a continuous sequence of activation across the population following gaze shifts that was evident in single trial spike rasters (Figure 2C and Supplemental Video 1). Applying this sorting to the mean PSTHs (Figure 2D) revealed a similar sequence, with positive peaks gradually becoming more delayed and an early negative component appearing (Figure 2D, left). We confirmed that this sequence was not an artifact of sorting by performing cross-validation, which showed that both the temporal order (Figure S2A) and response latencies (Figure S2B) were maintained when sorted on one half of the dataset (training set) and applied to the other half (test set; Pearson correlation coefficient, r=0.870, p=2.51e-140). The temporal sequence pattern was present, though weaker, in the non-preferred direction (Figure 2D, middle), and was absent for compensatory movements (Figure 2D, right). Notably, the four responsive groups generally maintained their grouping in the continuous sequence of temporal responses, with the early positive group showing the earliest peak positive response, followed by the late positive group, the biphasic group, and the negative group (Figure 2D, colorbar in left panel). Thus, the diverse responses to gaze shifts represent a temporal sequence across the population.

### Temporal dynamics of gaze shift responses depend on visual input

The selectivity of cells to gaze-shifting head and eye movements, but not compensatory movements that stabilize the gaze, led us to hypothesize that visual input may be driving the observed transients, as opposed to primarily motor efference or vestibular signals. To test the role of visual input in generating gaze-shift responses, we recorded neural activity during free movement both in the light and in complete darkness.

We clustered cells using the gaze shift PSTHs measured under normal light conditions (Figure 3A), and compared their responses in the light and dark (Figure 3B). The distinct cluster responses present in the light were absent in the dark, with most clusters (late positive, biphasic, and negative) showing an average suppression of activity with an onset before the start of the gaze shift (Figure 3C, n=265 units, 7 mice), consistent with previous reports of saccade suppression (Ibbotson and Krekelberg, 2011). As in the light, neurons also showed no response to compensatory movements in the dark (Figure 3D). Sorting units based on their response latency in the light revealed that the precise temporal sequence of responses found in the light was also not maintained in the dark (Figure 3E). Together with the absence of modulation around compensatory movements, this suggests that the dynamic responses following a gaze shift are due to the corresponding change in visual input.

**Figure 3.**
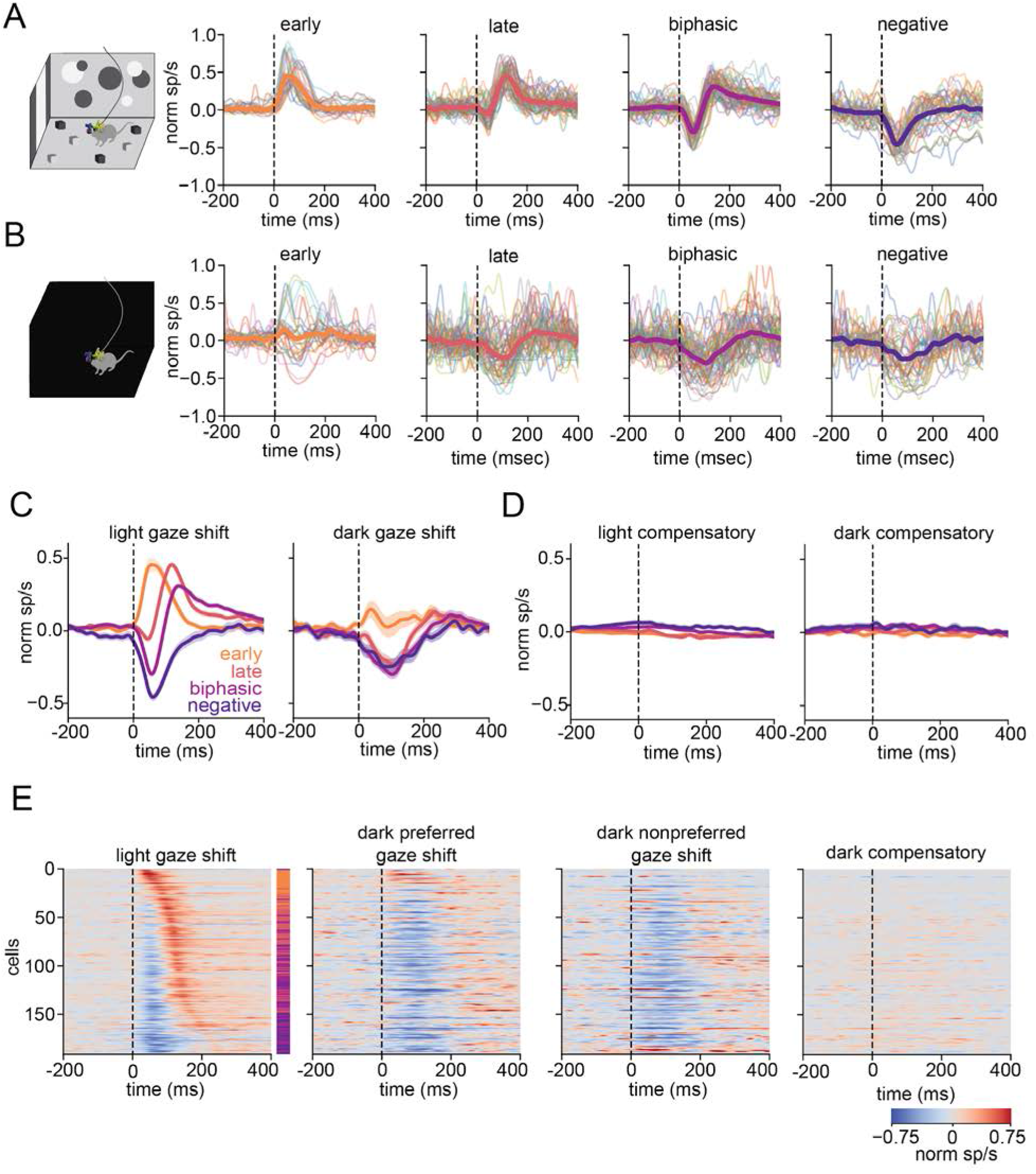
Temporal dynamics of gaze shift responses depend on visual input. (A) Normalized PSTHs in light condition for the preferred direction of gaze shifts, clustered using k-means weights used for cells in Figure 2. n=7 mice, 269 units (early n=27, late n=56, biphasic n=66, negative n=42, unresponsive=78). (B) Same as (A) for dark condition. Cells are grouped by their cluster from (A). (C) Mean gaze shift responses of each cluster in light (left) and dark (right) conditions with standard error. (D) Same as (C) for compensatory movements. (E) Temporal sequence of responses to gaze shifts and compensatory movements in the light and dark. All temporal sequences are sorted by the cell’s gaze shift response latency for the light condition (left). Cells unresponsive to eye movements in light conditions are shown at the bottom in shuffled order.

The mean response of the early positive group appeared flat in the dark, which resulted from a mix of units that showed suppression and a small fraction of units with strongly gaze direction-tuned responses that persisted in the dark (3.3% cells, n=9/269; Figure S3). The latter may represent a small population that receives a direct corollary discharge signal for saccades. However, for the vast majority of neurons, the rich diversity of responses we observe around gaze shifts in the light was replaced by a uniform suppression in the dark.

### Head-fixed flashed stimulus responses resemble freely moving gaze shift responses

We next sought to further test whether the observed responses to gaze shifts were visually driven by presenting stimuli that mimic the rapid onset of a new visual image that results from a gaze shift in a complex visual environment. Head-fixed mice were shown continuously flashed stimuli, including reversing checkerboard and sparse noise, prior to recording the same units during free exploration of the arena, and we compared the stimulus onset responses to gaze shift responses.

Time-aligned rasters and PSTHs for gaze shifts and the reversing checkerboard stimulus are shown for 3 example units in Figure 4A. Notably, the visually evoked response to continuously flashed full-field stimuli resembled the neuron’s response to gaze shifts, including positive, biphasic, and negative responses. This similarity was evident at the population level as well, as the mean stimulus onset PSTHs in response to the flashing checkerboard, clustered based on each unit’s gaze-shift response type, closely matched responses to gaze shifts (Figure 4B, left), as did mean responses to the shorter sparse noise stimulus (Figure 4B, right). To quantify the similarity between gaze-shift responses and the responses to flashed stimuli, we computed the correlation coefficient between PSTHs for gaze-shifts and the flashed stimuli for each unit (0-250 ms following onset). This revealed that gaze-shifting and flashed stimulus PSTHs were highly correlated for most units (Figure 4C).

**Figure 4.**
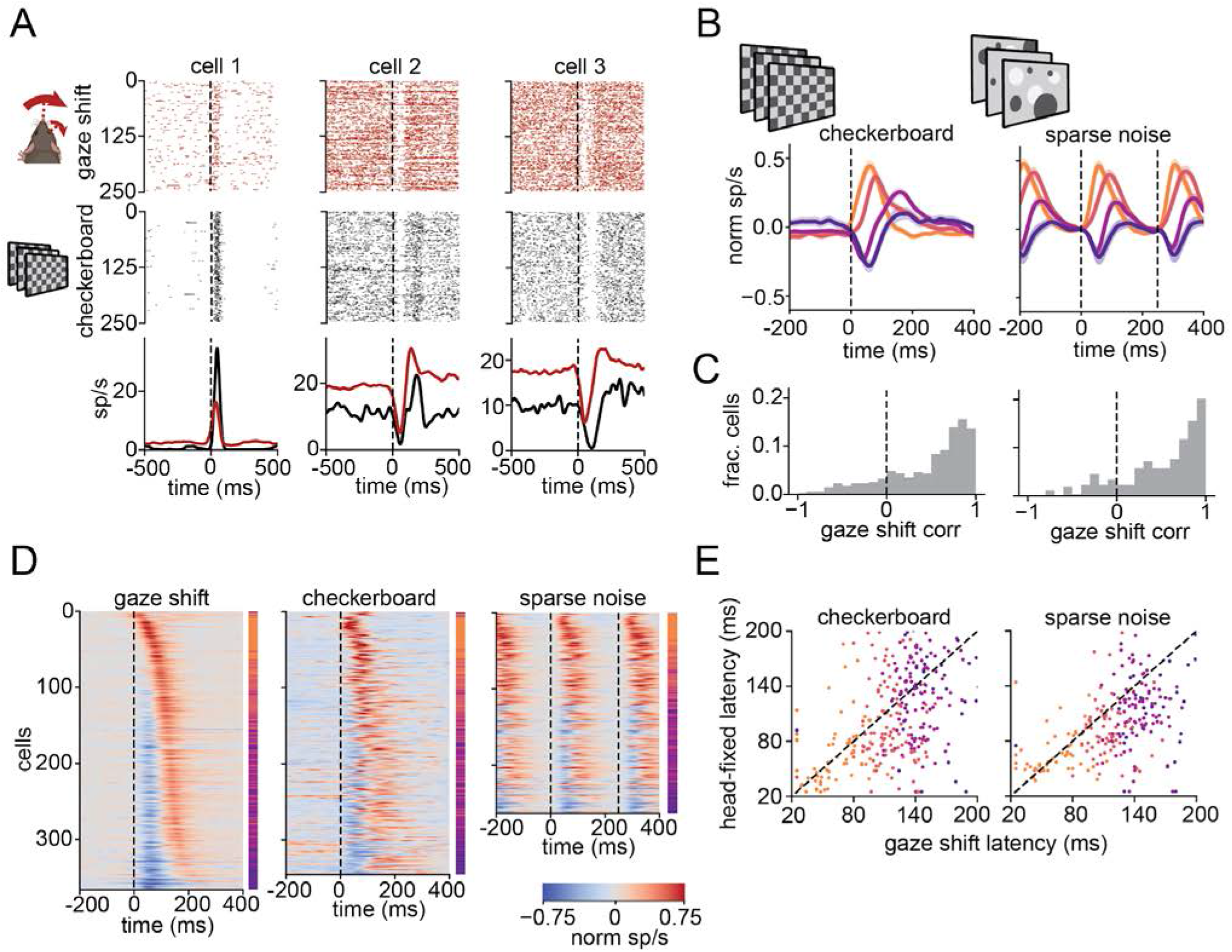
Head-fixed flashed stimulus responses resemble freely moving gaze shift responses. (A) Three example cells with positive (left), biphasic (middle) and negative (right) responses to freely moving gaze shifts. Spike rasters at the time of 250 randomly sampled gaze shifts during free movement (top) and 250 full-field reversals of a black and white checkerboard during a head-fixed recording of the same cell (middle). Bottom: gaze shift (red) and checkerboard (black) PSTHs for each cell. (B) Mean normalized response of cells to head-fixed flashed checkerboard (left) and sparse noise (right), clustered on their responses to freely moving gaze shifts. n=9 mice, 716 cells (checkerboard: responsive=472/716, early=67, late=100, biphasic=123, negative=55, unresponsive=127; sparse noise: responsive=333/716, early=51, late=69, biphasic=99, negative=44, unresponsive=70). (C) Pearson correlation coefficient between gaze shift PSTH and flashed checkerboard (left, mean=0.46, n=345) or sparse noise (right, mean=0.52, n=263). (D) Temporal sequence of gaze shift responses (left, includes cells responsive to gaze shifts and either flashed head-fixed stimulus, n=366), checkerboard (middle, cells responsive to both gaze shift and checkerboard, n=345) and sparse noise (right, cells responsive to both gaze shift and sparse noise, n=263) sorted on the latency of gaze shift responses in freely moving conditions. (E) Latency of positive peak in gaze-shifting responses versus head-fixed stimulus responses for checkerboard (left, r=0.487, p=5.89e-22) and sparse noise (right, r=0.526, p=4.19e-20).

Furthermore, the temporal sequence of checkerboard and sparse noise responses, when sorted based on each unit’s latency from freely moving gaze shifts, closely matched the temporal sequence from gaze shifts (Figure 4D). Correspondingly, the latency of each unit’s responses to head-fixed sequential stimuli was highly correlated with its latency for gaze-shift responses (Figure 4E). These results cannot be explained by the mouse performing gaze shifts with stimulus onset, as mice made only infrequent eye movements during head-fixation (sparse noise: 11.0 ± 10.1 saccades/min; checkerboard 13.6 ± 15.8 saccades/min). The close similarity between responses to gaze shifts during free movement and responses to sequentially flashed stimuli under head-fixed conditions strongly suggests that gaze shift transients primarily result from large, rapid changes to visual input.

### Spatial and temporal frequency tuning demonstrate coarse-to-fine processing around gaze shifts

We next sought to determine whether there was a relationship between the visual tuning properties of neurons and their temporal responses to gaze shifts. We presented drifting sinusoidal gratings of varying spatial and temporal frequencies to head-fixed mice prior to recording of the same units during free exploration of the arena. When organized by gaze shift response type, the mean PSTHs for gaze shift clusters to drifting gratings reveal a rapid response with clear onset transient in the positive clusters, and slower, more sustained responses in the biphasic and negative clusters (Figure 5A, B).

**Figure 5.**
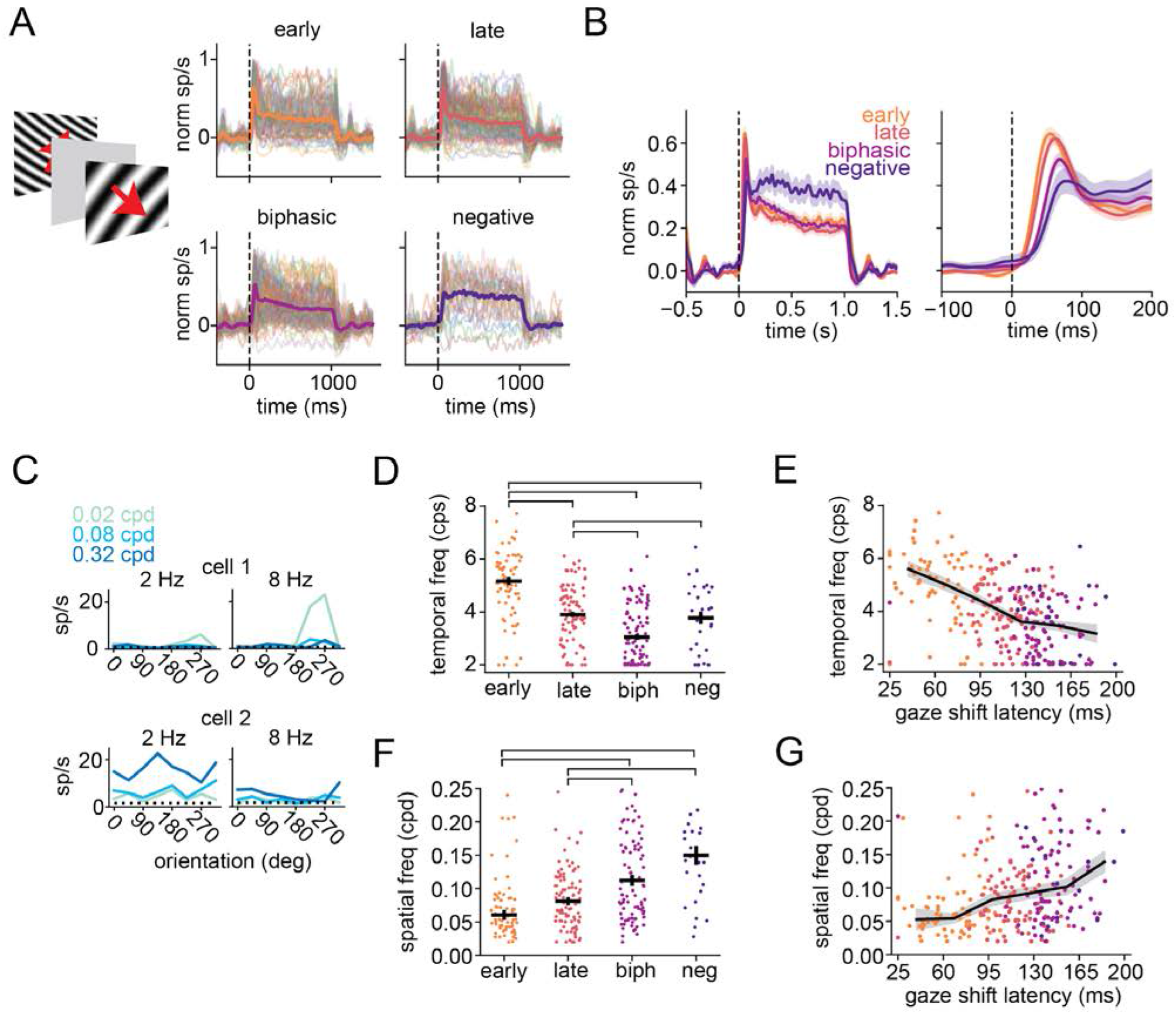
Spatial and temporal frequency tuning demonstrate coarse-to-fine processing around gaze shifts. (A) Head-fixed drifting gratings PSTHs for gaze shift response clusters. Stimulus is presented for 1 s with gray ISI between stimuli. n=384/716 responsive to gratings (early=71, late=96, biphasic=98, negative=29, unresponsive=90) with mean response overlayed. Cells below firing rate threshold are not shown. (B) Mean normalized gratings PSTHs clustered by gaze shift response for full stimulus presentation (left) and stimulus onset (right). (C) Orientation tuning curves for two example cells. Responses are shown for eight orientations of gratings (0 deg is the horizontal rightwards direction), three spatial frequencies, and low (left) and high (right) temporal frequencies) Spontaneous firing rates are shown as a horizontal dotted black line. (D) Preferred temporal frequency for gratings-responsive cells in each gaze shift response cluster, calculated as a weighted mean of responses. Median and standard error are shown for each cluster. Bars above indicate statistical significance at p<0.001, determined by pairwise ANOVA. (E) Preferred temporal frequency versus gaze shift response latency, for all cells responsive to gratings. Running median for all cells is overlaid. The color of each point indicates the cluster from gaze shift responses. (r=-0.468, p=2.12e-16). (F) Same as (D) for spatial frequency. 20/384 cells had a prefered spatial frequency greater than 0.25 cps are not visible in the scatter plot. These cells are included in the calculation of median. (G) Same as (E) for spatial frequency. (r=0.286, p=1.73e-6). Bars above indicate statistical significance at p<0.001, determined by pairwise ANOVA.

We computed the preferred temporal and spatial frequencies for each neuron based on its response to the drifting gratings (Figure 5C). The early positive group was tuned to the highest temporal frequency (TF), followed by late positive, then biphasic, and negative groups (Figure 5D). Correspondingly, the median TF to which units were tuned decreased as a function of increasing response latency (Figure 5E). The observed relationship between TF tuning and response latencies provides a potential explanation for the sequence we observe following gaze shifts: high TF-tuned cells respond quickly to gaze shifts, corresponding to the initial and abrupt change in visual input as a result of a shifting gaze. Low TF-tuned cells, conversely, respond later to gaze shifts during gaze fixation and maintenance, characterized by an absence of abrupt change to visual input.

The relationship between gaze-shift response and spatial frequency (SF) tuning was qualitatively opposite to TF tuning. The early positive group had the lowest SF preference, with increasing preferred SF across the late positive, biphasic, and negative groups (Figure 5F). Similarly, when sorting units based on gaze-shift response latency the median SF to which units are tuned increases as a function of latency (Figure 5G). Thus, the temporal sequence of responses following gaze shifts corresponds to a progression from low SF preferring neurons to high SF preferring neurons. These results are consistent with the coarse-to-fine theory of visual processing, which posits that the spatial components of new visual input are processed sequentially from low to high spatial frequencies (Hegdé, 2008).

### A coarse-to-fine temporal sequence in freely gazing marmosets

Given the finding of a coarse-to-fine visual processing sequence following gaze shift in freely moving mice, we wondered whether a similar temporal sequence occurs in V1 of non-human primates during active visual sampling. Mouse vision differs in several ways from non-human primates, but particularly striking is the lack of a fovea and high acuity vision (Huberman and Niell, 2011; Seabrook et al., 2017). We therefore recorded the activity of single units in foveal V1 of head-fixed marmosets to determine the neural response to saccades across complex visual scenes.

Marmosets were first head-fixed and allowed to freely view natural image stimuli (Figure 6A). Aligning spike times to the onset of saccades revealed gaze shift responses in marmoset that were similar to those in freely moving mice. Spike rasters and PSTHs from example units show positive, biphasic, and negative responses following saccades (Figure 6B). We clustered responsive units based on their normalized gaze shift PSTHs (k-means, k=4 as the cluster of non-responsive units was removed based on set criteria; Figure S4A). This resulted in qualitatively similar groups to those identified in mice (Figure 6C,D; Figure S4B,C). We also found a similar temporal sequence of responses across the marmoset V1 population after sorting by latency of peak positive response (Figure 6E), with the clusters maintaining their groupings across the temporal sequence in the same order as the mouse V1 units. We confirmed the sequence (Figure S2C) and latencies (Figure S2D) by cross validation.

**Figure 6.**
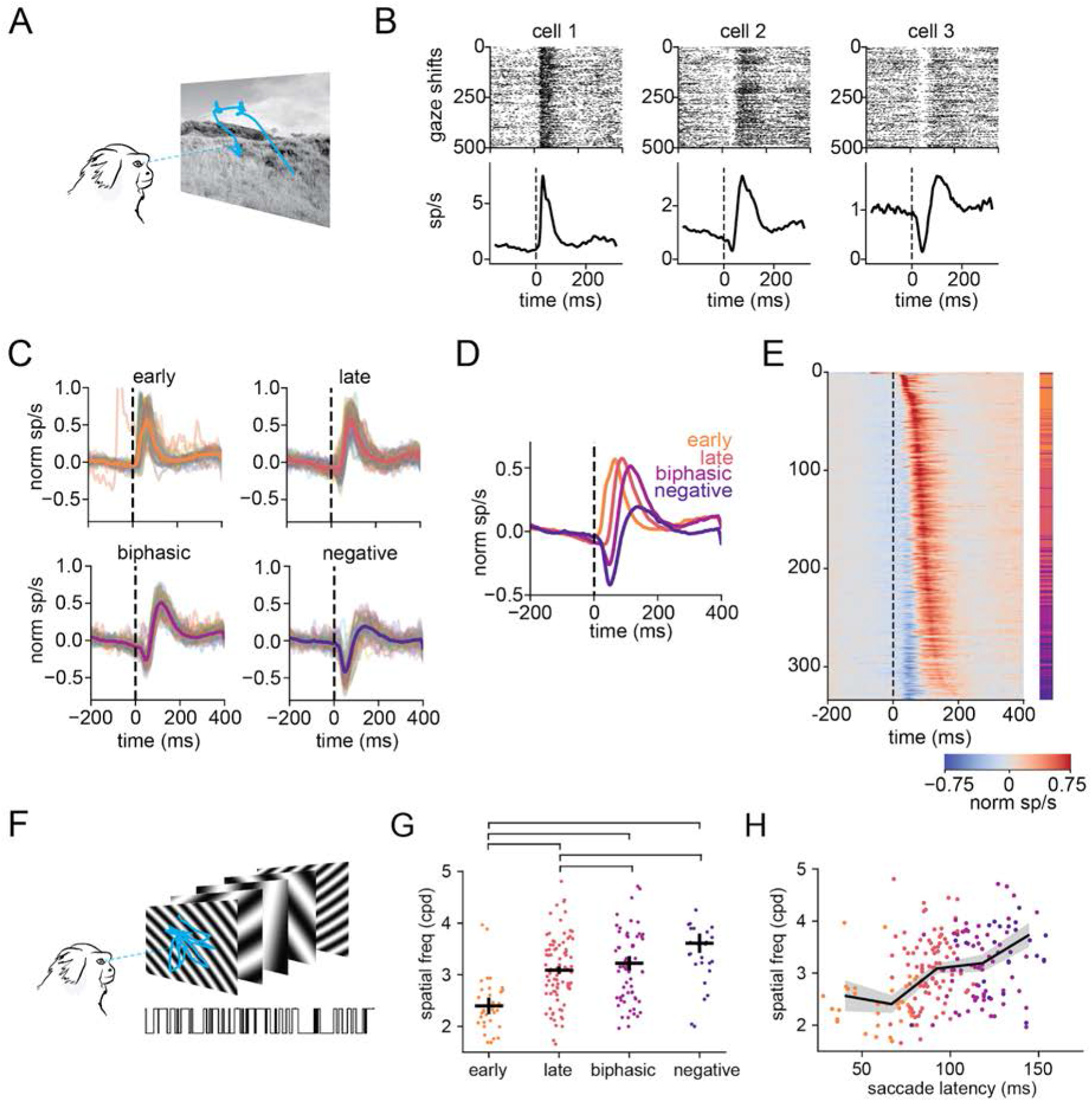
A coarse-to-fine temporal sequence in freely gazing marmosets. (A) Schematic of marmoset natural image free-viewing. (B) Three example cell’s spike raster (top) and PSTH (bottom) for 500 randomly selected saccades. (C) Normalized PSTHs for saccades, grouped by k-means clustering, with each cluster’s mean overlaid. n=2 marmosets, 334 units (early=64, late=136, biphasic=84, negative=50). (D) Mean normalized saccade response of cells in each cluster. (E) Normalized saccade PSTHs sorted along the y-axis by positive peak latency. Vertical colorbar indicates the response cluster for each cell using colormap from (C). (F) Schematic of grating stimulus. Sinusoidal gratings of varying spatial frequency and orientation were presented rapidly (60Hz) while monkeys performed saccades to forage for an embedded Gabor patch. (G) Preferred spatial frequency (SF) for each saccade response cluster, calculated as a weighted mean of responses for 4 presented SFs. Median and standard error are shown for each cluster. Only gratings-responsive cells are shown. n=238 units (early=41, late=88, biphasic=70, negative=39). (H) Preferred SF versus saccade response latency. Running median is overlaid. Point colors indicate response clusters. Outlier cells with a response latency below 25 ms (n=1) or above 160 ms (n=12) are not shown for plotting purposes, but included in statistical tests. n=334 units (early=40, late=88, biphasic=69, negative=29). (r=0.313, p=8.54e-7).

To determine whether this temporal sequence represents a similar coarse-to-fine mechanism, we examined SF tuning based on rapidly flashed sinusoidal gratings, presented while the monkey performed saccades to forage for embedded Gabor patches (Figure 6F; see Methods). This dataset was originally designed to study neural signals associated with visual foraging, but for the present study the same data provides an accurate estimate of SF tuning for the same neurons as in the recordings during natural image viewing. Specifically, the sinusoidal gratings represent a Hartley subspace that allows rapid mapping of receptive fields (Ringach et al., 1997) as well as orientation and SF tuning, though not TF tuning.

Similar to the SF tuning of the gaze shift response clusters in mice, marmoset neurons with earlier, positive saccade responses were tuned to low SFs, and biphasic/negative responses were tuned for higher SFs (Figure 6G). Likewise, we observed an increase in median preferred SF as a function of saccade response latency (Figure 6H). We thus find that marmoset foveal V1 shows a temporal sequence of gaze shift responses similar to that of freely moving mice, consistent with a coarse-to-fine pattern of visual processing shared across mouse and primate V1.

### Temporal tuning of neurons can explain diverse responses to gaze shifts

In both mice and marmosets, many units showed a decrease in activity immediately following a gaze shift (Figure 2,6), which was also present for flashed stimuli in the mouse data (Figure 4A,B). On the other hand, the PSTH of responses to drifting grating stimuli resembled visual responses that are typically observed in standard visual stimulus paradigms, with a positive response onset (Figure 5A,B). We hypothesized that this may be due to the presence of a gray inter-stimulus interval (ISI) in the grating stimuli, which is standard in presentation of many visual stimuli but is absent during gaze shifts in visually complex environments, where a new visual input arrives with each saccade without a return to baseline. Furthermore, the correspondence between temporal frequency preference and gaze-shift response, along with the similarity in response to flashed stimuli, suggests that these distinct responses could result from the neurons’ temporal dynamics to visual input changing in rapid succession.

To investigate this, we developed a simple linear-nonlinear model of a neuron’s response to a time-varying stimulus. We model the stimulus as a scalar input representing the visual drive that varies discretely across eight levels (such as eight different orientations of a grating stimulus) at 250 ms intervals (Figure 7A). We implemented two forms for this time-varying input – one in which the input returns to zero on alternate intervals, consistent with standard visual stimulus presentation with a gray ISI, and the other in which the input varies randomly at each interval without returning to zero, consistent with continuous stimulus presentation as occurs in sequential gaze shifts. We then modeled the neuron’s response as a linear temporal kernel applied to this stimulus, followed by an exponential nonlinearity to produce a spike rate. We used three temporal kernels: a biphasic kernel (Figure 7B) which acts as a differentiator and corresponds to transient, high temporal frequency responses; a unimodal positive kernel (Figure 7C) which acts as an integrator and corresponds to sustained, low frequency responses; and an intermediate between these two (Figure 7D).

**Figure 7.**
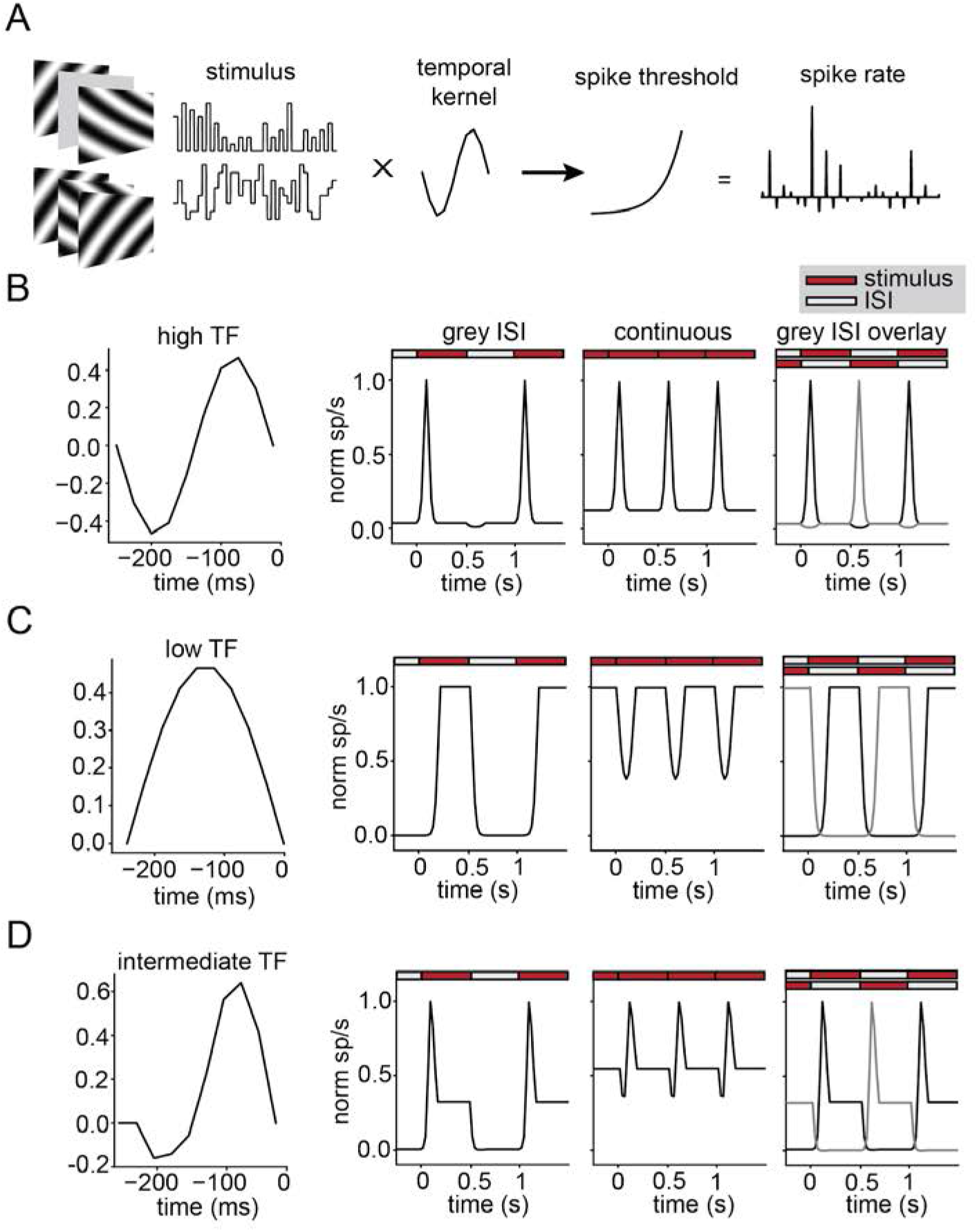
Temporal tuning of neurons can explain diverse responses to gaze shifts. (A) Schematic of modeling approach. A scalar stimulus either with (top left) or without (top right) an inter-stimulus interval (ISI) is passed through a variable temporal kernel and a non-linearity to generate a spiking output. (B) High temporal frequency (TF) kernel (left) and the resulting response to visual stimuli with an ISI (middle left) or continuously presented visual stimuli with no ISI (middle right). Responses to continuous stimuli are schematized as the simultaneous response to alternating stimuli with an ISI (right). (C) Same as (B) for low TF kernel. (D) Same as (B) for intermediate TF kernel.

Applying this model to the input with a gray ISI and calculating the mean PSTH across all stimulus transitions recapitulated expected responses from standard stimulus presentation paradigms, with the high TF kernel evoking a rapid transient response relative to ISI baseline, the low TF kernel evoking a slower but sustained response, and the intermediate kernel evoking a transient followed by sustained response. Strikingly, when the model was applied to the continuous input, it reproduced response types that we observed for gaze shifts, with the high TF kernel evoking short latency positive responses, the low TF kernel evoking negative responses, and the intermediate kernel evoking a biphasic response. The intuition behind this is that during continuous stimulus presentation, for the high TF kernel each new stimulus evokes a transient response and then returns to baseline, resulting in a positive peak. For the low TF kernel, rather than resting at a baseline firing rate before stimulus onset as would be the case with a gray ISI, the pre-stimulus rate is determined by the sustained response to the previous stimulus. Following the transition to a new stimulus, the firing rate drops due to the offset of the previous stimulus and the slow (low TF) response to the new stimulus, resulting in what appears as a negative response. In other words, the negative response need not result from active suppression, but simply the loss of response to the previous stimulus and lack of rapid onset response to the new stimulus. For the intermediate kernel, there is a combination of these two effects, leading to a brief drop from the previous stimulus offset followed by a mix of transient and sustained responses, giving a biphasic response. This intuition can be visualized by overlaying alternating responses to the gray ISI stimuli to approximate the continuous presentation condition (Figure 7B-D, far right), revealing the positive, biphasic, and negative patterns around stimulus transitions. Although this provides a visual intuition, it is important to note that the actual response during continuous presentation is a result of transitions across multiple levels of stimulus strength.

Together, this modeling suggests that the distinctive response patterns following a gaze shift can be explained, at least at the qualitative level, by each neuron’s temporal dynamics in response to individual stimuli, together with the fact that during gaze shifts, the response to each new stimulus rides on the response to the previous stimulus. Combined with the correspondence of spatial frequency preference with temporal dynamics, this gives rise to a sequence of processing across the oculomotor cycle, from the high TF/low SF response immediately following a gaze shift to the low TF/high SF response arising during the subsequent fixation.

## Discussion

We recorded the activity of V1 neurons as mice moved freely in an arena in order to determine how visual processing is influenced by eye and head movements at the cortical level. A wide scope of previous work has shown that activity of cortical neurons is correlated with movement, with V1 neurons showing responses to locomotion, head movements, and eye movements. We found a diverse array of responses to gaze-shifting, but not compensatory, head and eye movements in V1. When the animal moved in an arena lacking visual input (complete darkness), the diversity of gaze shift responses disappeared, and was replaced by a weak suppression of activity around the time of a gaze shift. Further, we found V1 neurons mimic their response to gaze shifts when shown full-field sequentially flashed stimuli under head fixation despite no corresponding head or eye movements. These findings suggest that the responses to gaze-shifting head and eye movements are in fact the result of changes to the visual input as opposed to motor efference or non-visual sensory reafferent signals. Additionally, we found neurons responding earlier to gaze shifts to be tuned for high temporal frequencies and low spatial frequencies, and neurons responding later to gaze shifts to be tuned for low temporal frequencies and high spatial frequencies. These findings are consistent with the coarse-to-fine model of visual processing, in which the visual system responds first to coarse aspects of the visual scene prior to finer details.

The temporal sequence that we observe arises from the ‘oculomotor cycle’ that occurs during natural vision, during which rapid saccadic eye movements are interspersed with slow fixational movements in a pattern that formats the visual input arriving on the retina (Boi et al., 2017). Our findings therefore reflect the fact that natural visual input is neither continuous nor organized into temporally-distinct ‘trials’ as in highly controlled experimental paradigms. This impacts both the statistics of the visual input and the corresponding neural dynamics, underscoring the need to study visual processing under naturalistic conditions.

These findings are remarkably consistent across mice and marmosets, both in terms of the specific temporal response profiles that we observe, and the correspondence with spatial frequency tuning. This is notable since although many basic aspects of cortical visual processing, such as orientation selectivity, are shared across mammals, one might expect that aspects related to active vision could differ. Primates have a pronounced foveal specialization accompanied by an expanded repertoire of eye movements, including voluntary targeted saccades independent of head movements. On the other hand, mice have less pronounced retinal specializations (van Beest et al., 2021; Bleckert et al., 2014), and eye movements are largely coupled to head movement and image stabilization, with little evidence of independently targeted saccades (Holmgren et al., 2021; Meyer et al., 2020; Michaiel et al., 2020; Zahler et al., 2021). Nonetheless, when analyzed in a shared framework of gaze shifts versus fixations, a similar computational principle emerges - a temporal sequence of response dynamics corresponding to coarse-to-fine processing.

### Temporal dynamics and tuning of gaze-shift responses

Our results provide new insight into the nature of neural signals measured around head and eye movements in natural conditions. In one of the few studies of free viewing of natural scenes in primates, a mix of positive and negative responses was observed following saccades (Gallant et al., 1998). Likewise, neural responses around microsaccades, when the fovea moved within the stimulus, revealed positive, negative, and biphasic responses (Leopold and Logothetis, 1998). Our results suggest that these diverse responses may correspond to the different temporal tuning of V1 neurons, which determines their response when the retinal input shifts abruptly following saccades or microsaccades. Likewise, a recent study observed modulation of V1 activity around head-orienting movements in rats (Guitchounts et al., 2020), with a diversity of positive and negative tuning in light and dark, with no systematic relationship. By measuring eye movements along with head movements, we found that most neurons primarily respond to the instants within an ongoing head movement that are associated with a saccadic movement of the eyes, which result in a gaze shift. Furthermore, our results explain the logic behind the diversity of head movement responses, based on the temporal sequence relative to gaze shifts and the tuning properties measured with grating stimuli - from early positive neurons with low SF / high TF tuning to biphasic and then negative neurons with high SF / low TF tuning (Figure 5,6, S1C). This time course also explains responses in the dark, as all but the earliest positive neurons shift to suppression in the dark (Figure 3).

Somewhat surprisingly, a large fraction of neurons had a negative response to gaze shifts in the light, as well as to flashed stimuli. Negative responses in visual neurons around the time of a saccade have long been hypothesized to be suppression of visual perception while performing an eye movement across the visual scene. We find that such negative responses are common, and represent part of a continuum from early positive responses through biphasic and finally largely negative responses. Notably, the neurons with the largest negative responses, and correspondingly latest positive responses, are also tuned to the lowest TFs. This suggests that the negative responses, at least in part, are a result of the neurons not responding to the high TF transient associated with the saccade, leading to a drop in activity, then slowly responding to the sustained visual input during fixation. This hypothesis is supported by a simple linear-nonlinear model, which is able to recapitulate the diverse temporal dynamics, including negative responses, based purely on different temporal kernels for a neuron’s response that correspond to high and low TF, or transient and sustained responses.

Notably, these temporal dynamics arise as a result of the abrupt arrival of new visual input, during both gaze shifts and continuously flashed stimuli, standing in stark contrast with the standard visual neuroscience approach of presenting a gray “‘ blank” during an inter-stimulus interval (ISI). ISIs allow neurons to return to their baseline firing rate in between visual stimuli in order to isolate responses with their corresponding stimulus; as a result, responses to stimuli that lack an ISI (e.g. change without interruption) can depend on the cell’s response to the previous input. Under natural conditions, however, visual stimulation is a continuous input without ISIs. As we found responses to rapidly shifting full-field stimuli in the head-fixed condition to recapitulate the responses to freely-moving gaze shifts, our findings lend support to the notion that neural dynamics depend on not only what stimuli are presented, but how they are presented. Accordingly, it is important to understand the visual stimulus as it arrives at the retina under natural conditions (Gibson, 1979).

Finally, the typical temporal response sequence around gaze shifts was replaced by universal suppression in the dark. This result is consistent with physiological saccadic suppression via corollary discharge, which has been reported in other species, including nonhuman primates and cats (Adey and Noda, 1973; Duffy and Burchfiel, 1975; Toyama et al., 1984). It is possible that this suppressive signal is also present in the light, but is largely masked by the strong visually driven responses. It may also partially contribute to the negative component observed in gaze-shifts in the light, amplifying the visually driven component. Alternatively, previous work in mice has also shown that responses to head rotation in V1 are luminance-dependent in a cell-type specific manner (Bouvier et al., 2020), and thus the sign of this signal may switch between the light and dark conditions. We also observed a small population of neurons (3.3%) that showed positive responses to gaze-shifting movements in both the light and dark. Similar results have been reported in head-fixed mice in the absence of visual input (Miura and Scanziani, 2021), though we observed a much smaller fraction of units with such responses.

### Coarse-to-fine processing during active vision

There is significant evidence from human psychophysics, using briefly presented stimuli, that supports coarse-to-fine processing, with low spatial frequency components being recognized before high spatial frequency components (Navon, 1977; Oliva and Schyns, 1997; Petras et al., 2019). At the neural level, single-unit recordings in head-fixed stimulus nonhuman primates and rodents have shown a progression of tuning from low spatial frequency to high spatial frequency following stimulus onset (Bredfeldt and Ringach, 2002; Mazer et al., 2002; Purushothaman et al., 2014; Skyberg et al., 2022). Here, we demonstrate that this coarse-to-fine visual mechanism is engaged during active vision in both mice and marmosets, resulting in a temporal sequence of responses to visual input with gaze shifts.

There are multiple reasons that have been proposed for why the visual system may utilize a coarse-to-fine order of processing (Hegdé, 2008). According to Marr’s primal sketch theory, initial visual processing may be devoted to setting a basic layout of the visual scene, followed by later activity to establish greater dimensionality and detail in the percept (Marr, 2010). Alternately, the order of responses may represent a matching to the typical input following a saccade and subsequent fixation, with lower SFs predominant immediately after a saccade due to the 1/f distribution of natural scenes, followed by a whitening of the distribution due to small movements during the fixations (Rucci et al., 2007). Additionally, recent work in rodents suggests that coarse-to-fine processing may support efficient coding of natural scenes (Skyberg et al., 2022).

The underlying mechanisms that generate coarse-to-fine processing in V1 are unknown. One possibility is the pooling of magnocellular and parvocellular type inputs from LGN in V1, which respond in both a temporally- and spatially-specific manner (Allen and Freeman, 2006). In primates, cells making up the magnocellular pathway convey information that is typically of high TF and low SF (‘ fast’ and ‘coarse’), while neurons of the parvocellular pathway are the opposite, showing tuning for information of low TF and high SF (‘ slow’ and ‘fine’). Although there isn’ t a clear delineation of magnocellular and parvocellular pathways in mice, a similar relationship between transient/sustained response types and corresponding SF tuning has been observed in mouse LGN (Piscopo et al., 2013), and previous work supports a between TF and SF tuning in mouse V1 (Gao et al., 2010). Alternatively, it has been proposed that a rapid refinement of SF tuning occurs as a result of recurrent processing in cortex, particularly resulting from active suppression of low SF responses (Bredfeldt and Ringach, 2002). Genetic tools available in mice may allow experiments to test these potential circuit mechanisms.

Together, these results present a dynamic view of processing during active vision, where each gaze shift initiates a temporal sequence supporting coarse-to-fine processing. This processing mode is shared across rodents and primates, and is dramatically different from the regime typically probed in head- and gaze-fixed vision. This emphasizes the importance of accounting for the actual dynamics of visual input that occurs during behavior under natural conditions, in order to understand the computations performed by the visual system.

## Author Contributions

PRLP and CMN conceived the project. PRLP and ESPL led mouse experiments. DMM and CMN led data analysis. NMC and SLS contributed to mouse experiments. ETTA contributed to data analysis. MCS generated audio track from mouse neural activity. JLY and JFM performed marmoset experiments. JLY, JFM, and DMM performed marmoset data analysis. All authors contributed to writing and editing the manuscript.

## Acknowledgements

We would like to thank Drs. Alex Huk, Cory Miller, David Leopold, Kasia Bieszczad, Michael Goard, and members of the Niell and Mitchell lab for helpful conversations and feedback on the manuscript. This work was supported by NIH grants UF1NS116377 (CMN and JFM), R01NS121919-01 (CMN), 4R00EY032179-03 (JLY), and R01EY030998-02 (JFM).

## Methods - Mouse

### Animals

All procedures were conducted in accordance with the guidelines of the National Institutes of Health and were approved by the University of Oregon Institutional Animal Care and Use Committee. Three-to eight-month old adult mice (C57BL/6J, Jackson Laboratories and bred in-house) were kept on a 12 h light/dark cycle. In total, 5 male and 6 female mice were used for this study.

### Surgery and Habituation

Mice were initially implanted with a titanium headplate over primary visual cortex to allow for head-fixation and attachment of head-mounted experimental hardware. After three days of recovery, widefield imaging (Wekselblatt et al., 2016) was performed to help target the electrophysiology implant to the approximate center of left monocular V1. A miniature connector (Mill-Max 853-93-100-10-001000) was secured to the headplate to allow repeated, reversible attachment of a camera arm, eye/world cameras and IMU (Michaiel et al., 2020; Parker et al., 2022). In order to simulate the weight of the real electrophysiology drive for habituation, a ‘dummy’ electrophysiology drive was glued to the headplate. Animals were handled by the experimenter for several days before surgical procedures, and subsequently habituated (∼45 min total) to the spherical treadmill and freely moving arena with hardware tethering attached for several days before experiments.

The electrophysiology implant was performed once animals moved comfortably in the arena. A craniotomy was performed over V1, and a linear silicon probe (64 or 128 channels, Diagnostic Biochips P64-3 or P128-6) mounted in a custom 3D-printed drive (Yuta Senzai, UCSF) was lowered into the brain using a stereotax to an approximate tip depth of 750 µm from the pial surface. The surface of the craniotomy was coated in artificial dura (Dow DOWSIL 3-4680) and the drive was secured to the headplate using light-curable dental acrylic (Unifast LC). A second craniotomy was performed above left frontal cortex, and a reference wire was inserted into the brain. The opening was coated with a small amount of sterile ophthalmic ointment before the wire was glued in place with cyanoacrylate. Animals recovered overnight and experiments began the following day.

### Hardware and recording

The camera arm was oriented approximately 90 deg to the right of the nose and included an eye-facing camera (iSecurity101 1000TVL NTSC, 30 fps interlaced), an infrared-LED to illuminate the eye (Chanzon, 3 mm diameter, 940 nm wavelength), a wide-angle camera oriented toward the mouse’s point of view (BETAFPV C01, 30 fps interlaced) and an inertial measurement unit acquiring three-axis gyroscope and accelerometer signals (Rosco Technologies; acquired 30 kHz, downsampled to 300 Hz and interpolated to camera data). Fine gauge wire (Cooner, 36 AWG, #CZ1174CLR) connected the IMU to its acquisition box, and each of the cameras to a USB video capture device (Pinnacle Dazzle or StarTech USB3HDCAP). A top-down camera (FLIR Blackfly USB3, 60 fps) recorded the mouse in the arena.

The electrophysiology headstage (built into the silicon probe package) was connected to an Open Ephys acquisition system via an ultra thin cable (Intan #C3216). Electrophysiology data were acquired at 30 kHz and bandpass filtered between 0.01 Hz and 7.5 kHz. We first used the Open Ephys GUI (https://github.com/open-ephys/plugin-GUI) to assess the quality of the electrophysiology data, then recordings were performed in Bonsai (Lopes et al., 2015) using custom workflows. System timestamps were collected for all hardware devices and later used to align data streams through interpolation.

During experiments, animals were first head-fixed on a spherical treadmill to permit measurement of visual tuning properties using traditional methods, then were transferred to an arena where they could freely explore. Recording duration was approximately 40 min head-fixed, and 1 h freely moving. For head-fixed experiments, a 70 cm monitor (BenQ GW2780) was placed approximately 27.5 cm from the mouse’s right eye, and visual stimuli were presented using Psychtoolbox-3 (Kleiner et al., 2007). Head-fixed stimuli were also recorded using the head-mounted world camera. First, we presented 15 min of a band-limited Gaussian noise stimulus (Niell and Stryker, 2008); spatial frequency spectrum 0.05 cpd to 0.12 cpd, flat temporal frequency spectrum with a low-pass cutoff at 4Hz) in order to confirm V1 targeting based on spike-triggered average receptive fields. We presented a contrast-reversing square-wave checkerboard stimulus for 2 min with a spatial frequency of 0.04 cpd and temporal frequency of 0.5 Hz. Flashed sparse noise consisted of full- and minimum-luminance circular spots on a gray background played for 5 min. Spots were 2, 4, 8, 16, and 32 deg in diameter and presented so that each size made up an equal fraction of the area on the screen, totaling 15% on average. Each stimulus frame was presented for 250 ms and immediately followed the previous frame with no ISI (Piscopo et al., 2013). Drifting sinusoidal gratings were presented at eight evenly-spaced directions of motion for three spatial frequencies (0.02, 0.08, 0.32 cpd) and two temporal frequencies (2, 8 cps) for 15 min, with a 1 s stimulus duration and 0.5 s gray ISI, and stimulus conditions randomly interleaved.

The arena was approximately 48 cm long by 37 cm wide by 30 cm high. A 61 cm monitor (BenQ GW2480) covered one wall of the arena, while the other three walls were clear acrylic covering custom wallpaper including black and white high- and low-spatial frequency gratings and white noise. A stimulus consisting of moving black and white spots (Piscopo et al., 2013) played continuously on the monitor while the mouse was in the arena. During recordings, the arena’s stimulus monitor played a sparse moving noise stimulus similar to the flashed sparse noise stimulus used during head-fixed recordings. For moving sparse noise, full- and minimum-luminance spots were 4, 8, and 16 deg in diameter. Each spot was assigned to move in one of eight evenly spaced directions and one of five speeds (10, 20, 40, 80, 160 deg/s). Spots appeared on the appropriate edge of the screen and moved across until they disappeared on the far edge. The floor was a gray silicone mat (Gartful) and was densely covered with black and white Legos to provide three-dimensional visual contrast. Small pieces of tortilla chips (Juanita’s) were lightly scattered around the arena to encourage foraging during the recording, however animals were not water or food restricted.

In order to eliminate all possible light within the arena during dark recordings, the entire behavioral enclosure was sealed in light-blocking material (Thorlabs BK5), all potential light sources within the enclosure were removed, and all external light sources were turned off. Animals were first recorded in the dark (∼20 min), then the arena lights and wall stimulus monitor were turned on (∼20 min). As a result of the dark conditions, the pupil dilated beyond the margins of the eyelids, which made eye tracking infeasible. Thus, prior to the experiment, one drop of 2% pilocarpine HCl ophthalmic solution was applied to the animal’s right eye to constrict the pupil to a size similar to that seen in the light. Once the pupil was restricted enough for tracking in the dark (∼3 min) the animal was moved into the dark arena for recording until the effects of the pilocarpine wore off (∼20 min), at which time the light recording began.

### Data preprocessing

The raw electrophysiology data from head-fixed and freely moving recordings acquired in the same session were concatenated into a single file for spike sorting, in order to allow us to confidently track single units across the entire experiment. Common-mode noise was removed by subtracting the median across all channels at each timepoint. Spike sorting was performed using Kilosort 2.5 (https://github.com/MouseLand/Kilosort), and isolated single units were then selected using Phy2 (https://github.com/cortex-lab/phy) based on parameters including contamination (<10%), firing rate (mean >0.5 Hz across entire recording), waveform shape, and autocorrelogram. After data were spike sorted in Kilosort and curated in Phy2, the resulting spike data were then split back out into individual recordings for analysis.

We extracted pupil position from the eye camera data as performed previously (Michaiel et al., 2020). Briefly, eye camera data were first deinterlaced to achieve 60 fps video, then eight points around the pupil were tracked with DeepLabCut (Mathis et al., 2018). We then fit an ellipse to these eight points and computed pupil position in terms of angular rotation. The head-mounted world camera was deinterlaced to achieve 60 fps video and distortions from the camera lens were corrected using OpenCV.

### Neural responses to head-fixed stimuli

To maintain internal consistency with eye cameras and obviate the need for a separate video stimulus synchronization signal, we determined stimulus onsets in head-fixed recordings directly from the head-mounted cameras. For the reversing checkerboard stimulus, we identified frame transitions from the head-mounted camera video based on k-means clustering (k=2) of the video into the two separate contrasts, and selected transitions between the clusters. For the flashed spare noise stimulus that updated every 250 ms, we determined the timestamps of stimulus onset based on the RMS pixel-wise change in the image, which showed clear peaks at stimulus transition. For drifting sinusoidal gratings, we determined direction of motion by computing optic flow from the worldcam video, spatial frequency based on the mean gradient magnitude from paired Sobel operators, and temporal frequency from the mean Fourier transform of each pixel over time. Neuronal tuning was based on the evoked firing rate for each stimulus condition, computed as the mean rate from 25 to 1000 ms following stimulus onset, minus the mean baseline rate in the 500 ms before stimulus onset. Each cell’s preferred temporal frequency was determined using a weighted mean of evoked responses for each temporal frequency at the cell’s preferred spatial frequency and orientation, and likewise for spatial frequency at the cell’s preferred temporal frequency and orientation.

### Neural responses to eye/head movements

Horizontal head rotation velocity was extracted from the IMU and converted to deg/sec, then interpolated to eye-camera frame timestamps. Frames were considered the timepoints of a head movement if there was a head velocity >60 deg/s in the leftward or rightward direction. We defined leftward and rightward directions as the direction of head movement from the animal’s perspective. Eye/head movements were separated into either gaze-shifting or compensatory movements using gaze velocity, defined as the sum of horizontal eye and head velocities (Michaiel et al., 2020). Head-movements resulting in a high gaze velocity (>240 deg/s) were considered gaze-shifting, while those resulting in a low gaze velocity (<120 deg/s) were considered compensatory. For eye/head movements spanning multiple eye camera frames, only the first time point, representing the onset of the movement, was used. Compensatory movements which occurred 250 ms before or after a gaze-shifting movement were excluded to avoid contamination by gaze shifts. The same criteria were used to identify saccades during head-fixed recordings, but based on eye velocity alone (>240 deg/s) since the head was immobilized.

Neural responses around eye/head movements and flashed stimuli were calculated as a peristimulus time histogram (PSTH) from electrophysiology spike times using kernel density estimation with a gaussian kernel with bandwidth of 10 ms, sampled at 1 ms intervals. Gaze shift direction selectivity index was calculated as the difference between the preferred and non-preferred maximum modulations divided by their sum. To normalize PSTHs, we computed the baseline (R_b_) as the mean firing rate -1000 to -200 ms before the event. Because the head-fixed sparse noise stimulus was presented every 250 ms, we used the spike rate at the time of the stimulus onset, 0 ms, as the baseline firing rate to avoid including responses to previously presented flashed stimuli in the calculation of R_b_. We subtracted this baseline to give an evoked firing rate, and then calculated modulation as the maximum absolute value of the evoked rate within a response range of -250 ms before to 250 ms after the event. We used the horizontal head movement direction with the higher gaze shift modulation as the unit’s preferred direction. When normalizing the PSTH for an eye/head movement, we used max(R) from gaze shifts in the prefered direction so that all eye/head responses are normalized relative to the unit’s best response.

Units were considered responsive to gaze shifts if they were modulated by the onset of a gaze shift in its preferred horizontal head direction by >1 sp/s and showed a 10% modulation of its normalized spike rate in the 250 ms following the movement. To cluster units with similar gaze shift responses, we first performed Principle Component Analysis (PCA) on the normalized gaze shift responses in each unit’s prefered direction in the period from -50 to 300 ms relative to the gaze shift onset. Units which did not meet the above spike rate criteria had their input for PCA replaced with an array of zeros to exclude their non-responsive PSTHs from clustering, while seeding the formation of an unresponsive group of units for clustering. We used the first 4 principal components, which cumulatively explained 95% of the variance, to perform k-means clustering (k=5), resulting in 4 clusters of responsive units and 1 cluster of unresponsive units. To maintain consistency, the same PC weights and k-means model were applied to cluster the smaller number of units that were acquired subsequently in light/dark experiments.

The latency of a neural response was calculated as the maximum of its PSTH in the period between 25 to 250 ms relative to the onset of the event. Responses were sorted by the latency of their peak for the preferred direction of gaze shifting eye/head movements. Peak latency was not calculated for units unresponsive to gaze shifts. Peak latency sorting was cross-validated by randomly assigning gaze shifts into a train and test set, calculating a PSTH for each half of the data, and sorting the test set by the peak times of the training set (Figure S2).

To identify units that respond positively in dark conditions, we selected units which were positively modulated in the dark by the onset of a gaze shift by >1 sp/s and 10% of their normalized spike rate in the 100ms following the gaze shift onset.

To group units into narrow- and broad-spiking putative cell types, we first normalized the mean spike waveform of each unit as W_norm_ = (W - W_b_) / max(abs(W - W_b_)). We performed k-means clustering on the normalized waveforms (k=2). The resulting clusters closely resembled the segregation of narrow and broad spiking units in previous studies based on explicit waveform criteria (Niell and Stryker, 2008).

Laminar depth was calculated using the multi-unit LFP power of all sites on each single shank of the linear silicon probes, with the peak of LFP power considered the center of layer 5 based on (Senzai et al., 2019).

## Methods - Marmoset

### Animals and Surgical Procedures

All surgical and experimental procedures were approved by the Institutional Animal Care and Use Committee at the University of Rochester in accordance with the US National Institutes of Health guidelines. Two adult male common marmosets (*Calithrix jacchus*) aged 2 and 7 years were used in this study. Marmosets underwent an initial surgery to implant a headpost to stabilize their head during behavioral sessions as described previously (Mitchell et al., 2014; Nummela et al., 2017). A second surgery was performed to implant a recording chamber with a 3×3 mm craniotomy over foveal and parafoveal V1. The exposed dura was sealed under a thin layer (< 1 mm) of a silicone elastomer (Spitler and Gothard, 2008).

### Hardware and Recordings

Electrophysiological recordings were performed using 2×32 channel silicon electrode arrays (http://www.neuronexus.com). Probes included 2 sharpened tip shanks of 50µm width spaced 200 µm apart, each containing 32 channels separated by 35 µm. In one animal we used a semi-chronic microdrive (EDDS Microdrive system, https://microprobes.com) to place electrodes in cortex for 1-2 weeks over which we made 3-6 recordings. In the second animal we used a custom micro-drive (https://marmolab.bcs.rochester.edu/resources/) to place and remove electrodes daily. Arrays were lowered slowly through silastic into cortex using a thumb screw.

Data were amplified and digitized at 30 kHz with Intan headstages (Intan) using the Open Ephys GUI (https://github.com/open-ephys/plugin-GUI). The wideband signal was high-pass filtered by the headstage at 0.1 Hz, preprocessed by common-average referencing across all channels, and then high-pass filtered at 300 Hz. The resulting traces were spike sorted using Kilosort2. Outputs from the spike sorting algorithms were manually labeled using ‘phy’ GUI (https://github.com/kwikteam/phy). Any units that were either physiologically implausible based on the lack of a waveform with a trough followed by a peak or with an inter-spike interval (ISI) distribution with more than 1% of the spikes under 1 ms were excluded from analyses.

### Eye-tracking and saccade detection

Gaze position was monitored using infra-red eye tracking methods described previously (Yates et al., 2022) Briefly, the 1^st^ and 4^th^ Purkinje images (P1 and P4) were visualized using a collimated IR light source and tracked at 593 frames per second to estimate the 2D eye angle. The eye tracker was manually calibrated to adjust the offset and gain (horizontal and vertical) by showing marmoset monkeys small windowed face images at different screen positions to obtain their fixation as described previously (Mitchell et al., 2014; Nummela et al., 2017). Saccadic eye movements were identified automatically using a combination of velocity and acceleration thresholds (Cloherty et al., 2020).

### Head-fixed visual stimuli and free-viewing tasks

Visual stimuli were presented on a Propixx Projector (Vpixx) with a linear gamma table using Psychtoolbox-3 (Kleiner et al., 2007) and Matlab (Mathworks). Stimulus and physiology clocks were aligned using a Datapixx (Vpixx) following the methods previously described (Eastman and Huk, 2012). Stimulus code is available at (https://github.com/jcbyts/MarmoV5).

Marmosets performed free-viewing and visual foraging tasks. For natural image stimuli, the marmosets were allowed to free-view a screen containing a full-field (± 15 visual degrees) grayscale natural image for 10 s and then received a single drop of juice (5-10 µl of marshmallow water). On the foraging trials the marmosets viewed a random sequence of flashed full-field gratings (60 or 120 Hz frame update of varying orientation and spatial frequency gratings) as they searched for a small Gabor target (Yates et al., 2022). A drop of juice reward was given for fixating within 2 degrees of the small target for more than 100 ms after which time it was repositioned to another random location with 5 visual degrees of the screen center. During foraging the full-field flashed gratings were presented at 25% contrast and were drawn from a polar grid of 8 equally spaced orientations and a grid of 4 log spaced range of spatial frequencies from either 1 to 8 cycles/deg or 2 to 16 cycles/deg (8×4 stimulus space). On each video frame either a single grating was presented or with 50% chance a blank gray background was shown.

### Estimation of orientation and spatial frequency tuning

To measure orientation and spatial frequency tuning of individual neurons we analyzed the response to flashed full-field sine-wave gratings described above for the foraging task. A linear estimate of the receptive field can be recovered from this stimulus using subspace reverse correlation by correlating the firing rate during each video frame with the stimulus history (Ringach et al., 1997). We first computed a temporal kernel of the response to gratings as compared to blanks. The average temporal response at latencies from -50 to 200 ms (-3 to 12 video frames at 60 hz) was computed by time-locking on video frames including gratings and averaging. It was compared against the baseline firing rate, which was estimated by instead time-locking to blank video frames. If a peak was identified in the temporal response between 20 to 80 ms that was above 8 standard deviations above the baseline then the neuron was included in analyses for tuning. We marked a temporal window around the peak of the temporal kernel for which the response was at adjacent frames connected to the peak were also significant (5 standard deviations above baseline) including frames connected to the peak.

To estimate orientation and spatial frequency tuning the mean response was computed from the identified temporal window for each grating stimulus. The joint tuning for orientation and spatial frequency were defined by the mean responses in the 8×4 grid of stimuli sampled from which we averaged across orientation or spatial frequency to get the marginal tuning curves. To determine if neurons exhibited a selective response we computed a tuning index by taking the standard deviation of the values in the tuning curve divided by the mean. Neurons with weak tuning, having an index less than 0.2 in both dimensions, were not included for subsequent analyses that aimed to correlate tuning preference with saccade modulation. The preferred tuning was computed using a weighted mean from the tuning curve instead of taking the peak at a discrete value. To emphasize the peak in the weighted mean we used a weighting function computed by subtracting the mean of the tuning curve with the resulting negative values floored at zero and then squaring the remaining non-zero values.

### Neural responses to eye movements

Neural responses around eye movements were computed from -200 ms before to 400 ms after the saccadic onset. The peristimulus time histogram (PSTH) was computed using kernel density estimation with a Gaussian kernel with half-bandwidth of 10 ms, sampled at 1 ms intervals. To normalize PSTHs (R), we computed the baseline firing rate (R_b_) from -100 to 0 ms before saccade onset and normalized the PSTH as R_norm_ = (R - R_b_) / max(R). The maximum of R was computed from 0 to 400 ms. The PCA was computed on the normalized PSTH from -200 to 400 ms. The 6 PCs explained 95% of the variance and were k-means clustered (k=4) to group units into saccade response types.

### Statistical Analysis

Statistical tests across cell types were performed using ANOVA with correction for multiple comparisons. Correlations between latencies and response properties were tested using Spearman’s correlation coefficient. Bar plots show mean and standard error of the mean unless otherwise noted

## Supplemental Figures

**Figure S1.**
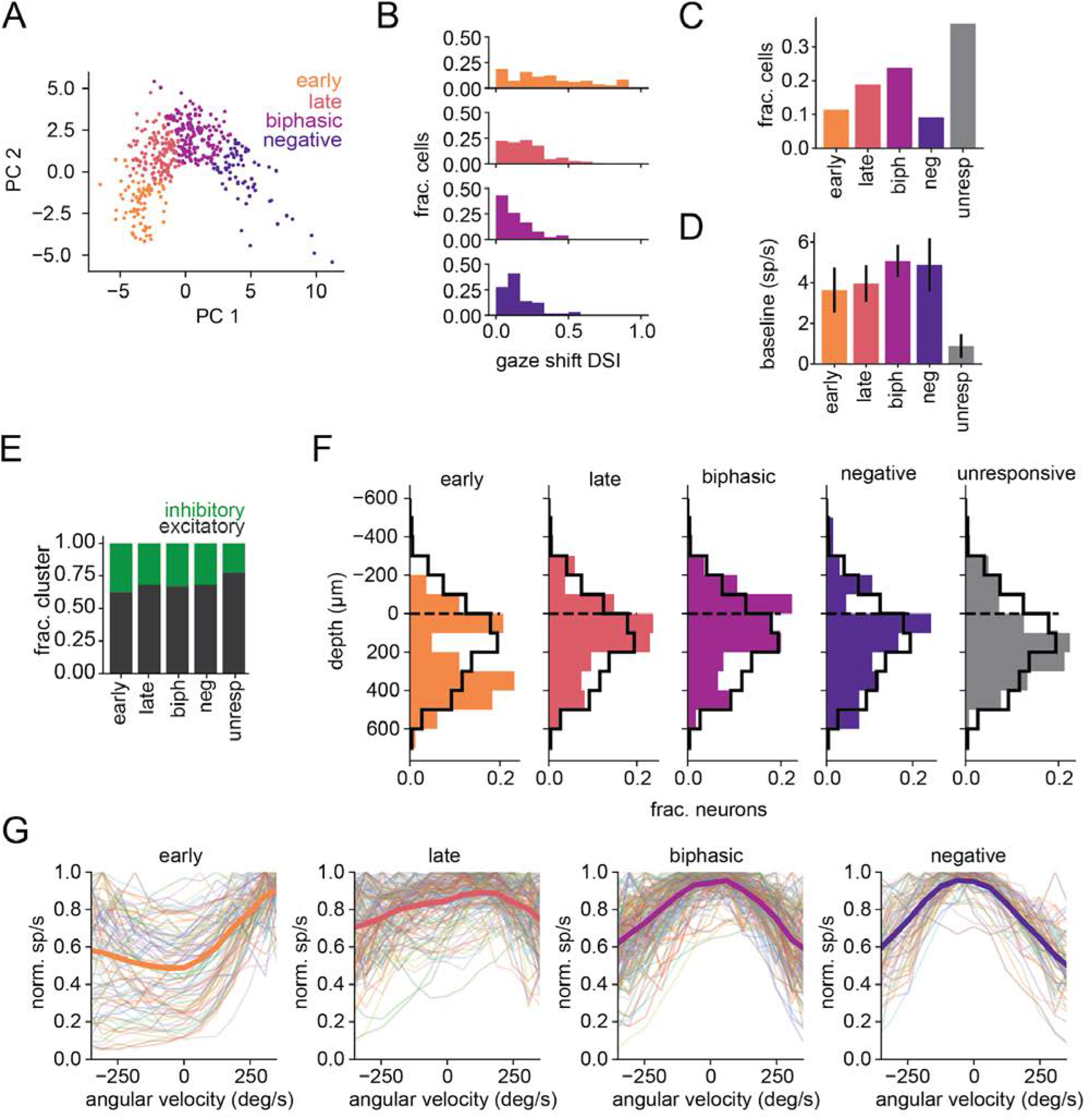
Additional characterization of gaze shift response types. (A) PCA of gaze shift PSTHs. Only the two PCs with the highest explained variance are shown. Cells in the scatter plot are colored by the cluster they were assigned by k-means clustering of PCs. (B) Gaze shift left/right direction selectivity index by cluster. (C) Fraction of units in each gaze shift response cluster. (D) Median baseline firing rate of units during freely moving recordings. (E) Putative cell type for each gaze-shifting response cluster. Excitatory and inhibitory groups were identified by k-means clustering on spike waveform. (F) Laminar depth of all cells, determined using the local field potential from multi-unit activity power along each shank of the probe. Black outline shows the distribution of depths for all cells. Dashed line (0 μm) is the estimated depth of cortical layer 5, to which depths were aligned. (G) Normalized horizontal angular velocity tuning for all cells, separated by response clusters. Positive values for angular velocity represent each unit’s preferred horizontal direction of gaze shift.

**Figure S2.**
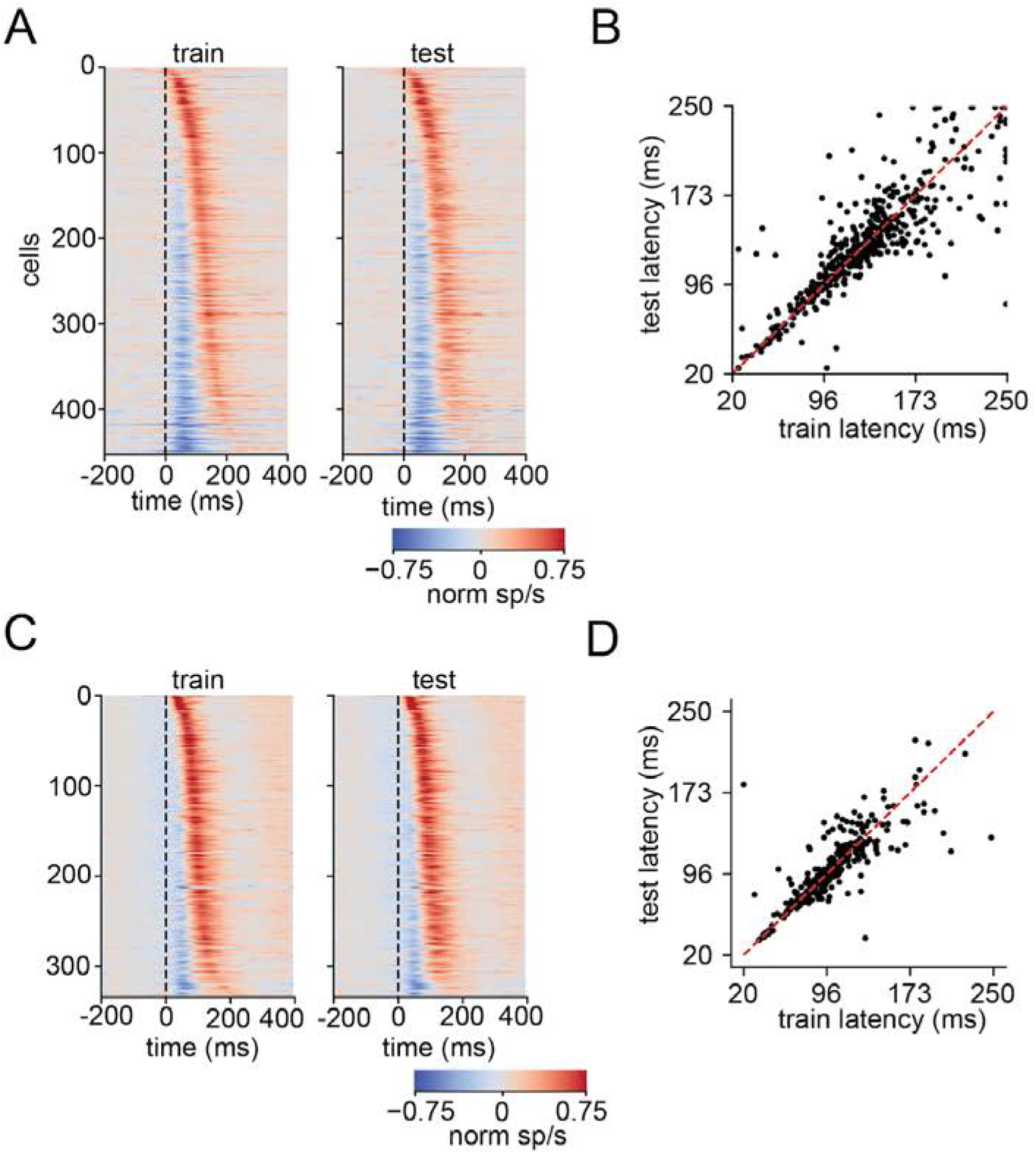
Cross validation of response latencies. (A) Cross-validation for mouse gaze shift PSTHs of all responsive cells. Gaze shift times were randomly divided into two sets used to calculate PSTHs in the train (left) and test (right) sets. The test set was sorted by the latency of positive peak in the train set. (B) Latency of gaze shift response for train versus test sets (r=0.870, p=2.51e-140). (C) Same as (A) for marmoset saccades. (D) Same as (B) for marmoset saccades (r=0.875, p=1.44e-106).

**Figure S3.**
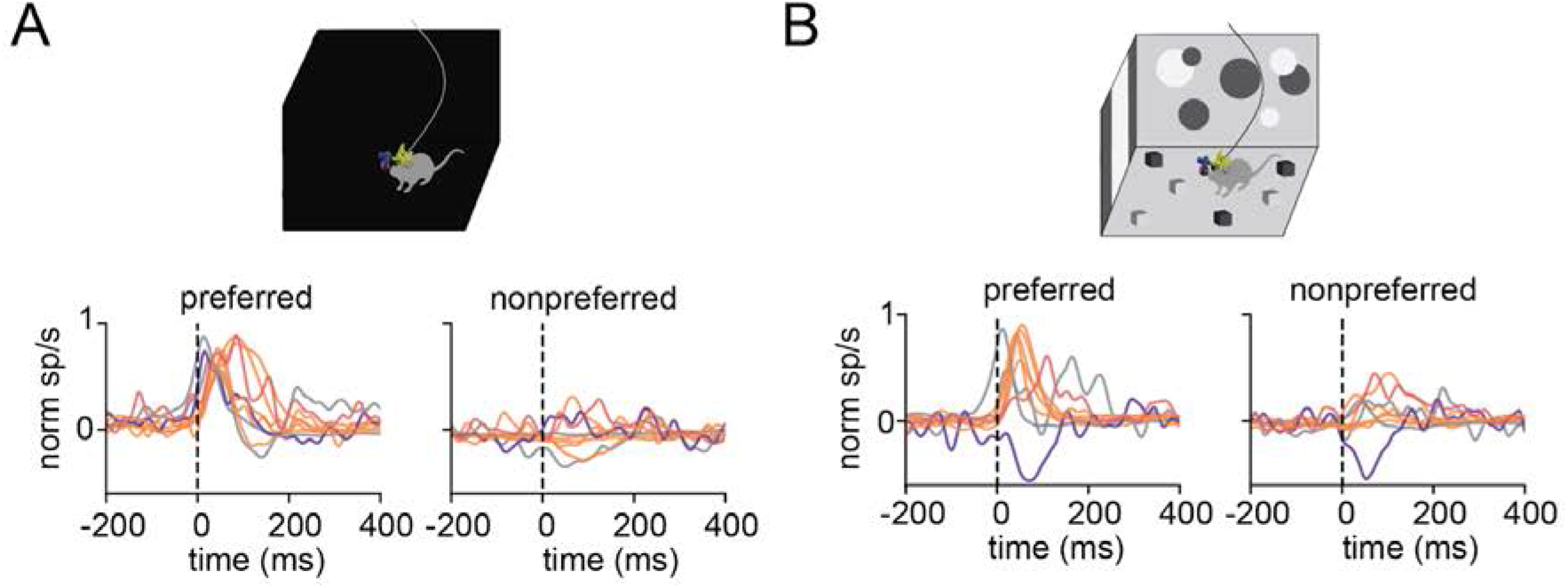
Additional characterization of responses in the dark. (A) Dark condition PSTHs for cells that responded to gaze shifts in freely moving dark conditions. Units are colored by clustering from responses in light condition. n=9/269 (early=5, late=1, biphasic=0, negative=1, unresponsive=2). (B) Light condition PSTHs for cells in (A).

**Figure S4.**
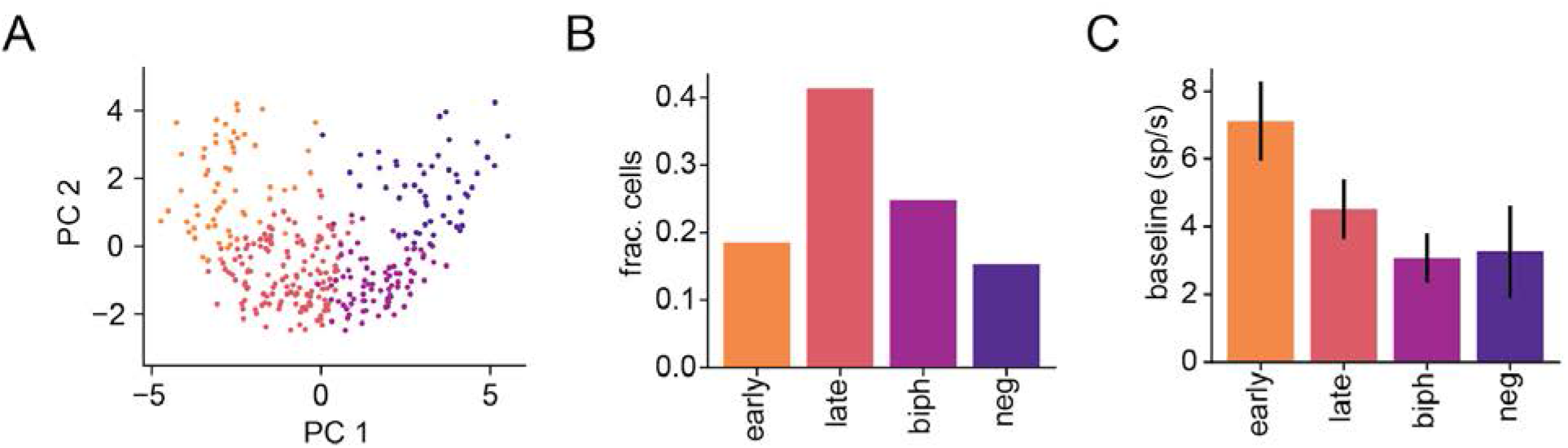
Additional characterization of marmoset saccade responses. (A) PCA of marmoset gaze shift PSTHs for the 2 PCs with highest explained variance, colored by k-means clusters. (B) Fraction of units in each saccade response cluster. (C) Median baseline firing rate of units in each cluster.

## Supplemental Videos

**Video S1:** A temporal sequence across the population following gaze shifts. Corresponds to Figure 2C. Experimental data from a 3 s period of freely moving activity. Top left: eye camera video. Top right: estimated visual input, based on world camera video corrected for eye position as in (Parker et al., 2022). Bottom: spike rasters for simultaneously recorded gaze shift-responsive units (n=99), color-coded by gaze-shift cluster and ordered from short to long gaze shift response latency along the y axis. Black arrows above the spike rasters indicate the time of gaze shifts. In the audio channel, individual spikes from 35 units are represented by notes mapped into pitch based on the temporal sorting, from short latency as low pitch to long latency as high pitch. Saccade times are represented as percussive notes.

